# A novel zebrafish model of *SPEG*-related centronuclear myopathy (CNM): characterization and comparison with other CNM model zebrafish

**DOI:** 10.1101/2021.12.22.473918

**Authors:** Karla G. Espinosa, Salma Geissah, Linda Groom, Jonathan Volaptti, Ian C. Scott, Robert T. Dirksen, Mo Zhao, James J. Dowling

## Abstract

Centronuclear myopathy (CNM) is a congenital neuromuscular disorder caused by pathogenic variation in genes associated with membrane trafficking and excitation-contraction coupling (ECC). Bi-allelic autosomal recessive mutations in striated muscle enriched protein kinase (*SPEG*) account for a subset of CNM patients. Previous research has been limited by the perinatal lethality of *Speg* knockout mice. Thus, the precise biological role of SPEG in skeletal muscle remains unknown. To address this issue, we generated zebrafish *spega, spegb*, and *spega/spegb* (*speg*-DKO) mutant lines. We demonstrate that *speg*-DKO zebrafish faithfully recapitulate multiple phenotypes associated with human CNM, including disruption of the ECC protein machinery, dysregulation of calcium homeostasis during ECC, and impairment of muscle performance. Taking advantage of the availability of zebrafish models of multiple CNM genetic subtypes, we compared novel and known disease markers in *speg*-DKO with *mtm1*-KO and *DNM2*-S619L transgenic zebrafish. We observed desmin (DES) accumulation common to all CNM subtypes, and DNM2 upregulation in muscle of both *speg*-DKO and *mtm1*-KO zebrafish. In all, we establish a new model of *SPEG*-related CNM, and identify abnormalities in this model suitable for defining disease pathomechanisms and evaluating potential therapies.

**Summary Statement:** We created a novel zebrafish *speg* mutant model of centronuclear myopathy that recapitulates key features of the human disorder and provides insight into pathomechanisms of the disease.

## Introduction

Congenital myopathies are neuromuscular disorders that typically present at birth with hypotonia and weakness (Louis et al., 2022). The prevalence of congenital myopathies is at least 1 in 26,000 (Amburgey et al., 2011). A common congenital myopathy subtype is centronuclear myopathy (CNM), which is defined pathologically by perinuclear organelle disorganization and increased centralized myofibre nuclei, and is clinically associated with respiratory failure, wheelchair dependence, and early mortality (Nance et al., 2012, Jungbluth et al., 2008). CNM is a genetic disease, caused by pathogenic variants in at least five genes: X-linked recessive CNM (XLMTM) caused by *MTM1* (myotubularin 1) (Laporte et al., 1996), autosomal dominant CNM caused by *DNM2* (dynamin 2) (Bitoun et al., 2005), and autosomal recessive forms of CNM caused by *BIN1* (bridging integrator 1) (Nicot et al., 2007a), *RYR1* (ryanodine receptor 1) (Wilmshurst et al., 2010), and *SPEG* (striated muscle enriched protein kinase) (Agrawal et al., 2014b).

Most mutations reported in CNM patients are associated with primary or secondary defects of the excitation-contraction coupling (ECC) machinery (Gonorazky et al., 2018, Tasfaout et al., 2018, Jungbluth et al., 2008, Dowling et al., 2021). In skeletal muscle, ECC occurs at triads, which consist of centrally located sarcolemmal invaginations called transverse tubules (or T-tubules) flanked on either side by terminal cisternae of the sarcoplasmic membranes (or tSRs) (Fig. 1A). These junctional membranes are connected by a mechanical interaction between the CaV1.1 subunit of the dihydropyridine receptor (DHPR) in the T-tubule membrane (with additional essential proteins including STAC3 and the DHPR β1a subunit) and the ryanodine receptor 1 (RyR1) Ca^2+^ release channels in the tSR. ECC is initiated when an action potential propagates down the T-tubule to cause voltage-driven conformational changes in DHPR, which then trigger activation of RyR1 to release Ca^2+^ stored in the tSR. The resulting surge in myoplasmic Ca^2+^ promotes actin-myosin cross bridging and sarcomere shortening (i.e. muscle contraction). Notably, although the DHPR/RyR1 interaction is essential for ECC function, the junctional membranes of the triad develop independently to the recruitment of DHPR or RyR1 to the region [as reviewed by (Al-Qusairi and Laporte, 2011)]. The tSR membrane also contains structural/regulatory proteins such as triadin, junctin, and calsequestrin (Zhang et al., 1997, Park et al., 2004), as well as other regulatory proteins that modulate RyR1 channel activity and maintain RyR1 integrity (Zhang et al., 1997, Costello et al., 1986, Treves et al., 2009, Ríos and Györke, 2009, Guo and Campbell, 1995, Wium et al., 2016, Caswell et al., 1999, Groh et al., 1999, Goonasekera et al., 2007). While many of the molecular components of the triad are known, the precise molecular mechanisms that underlie triad formation and maintenance, as well as triad disruptions in CNM, remain largely elusive.

**Figure 1.**
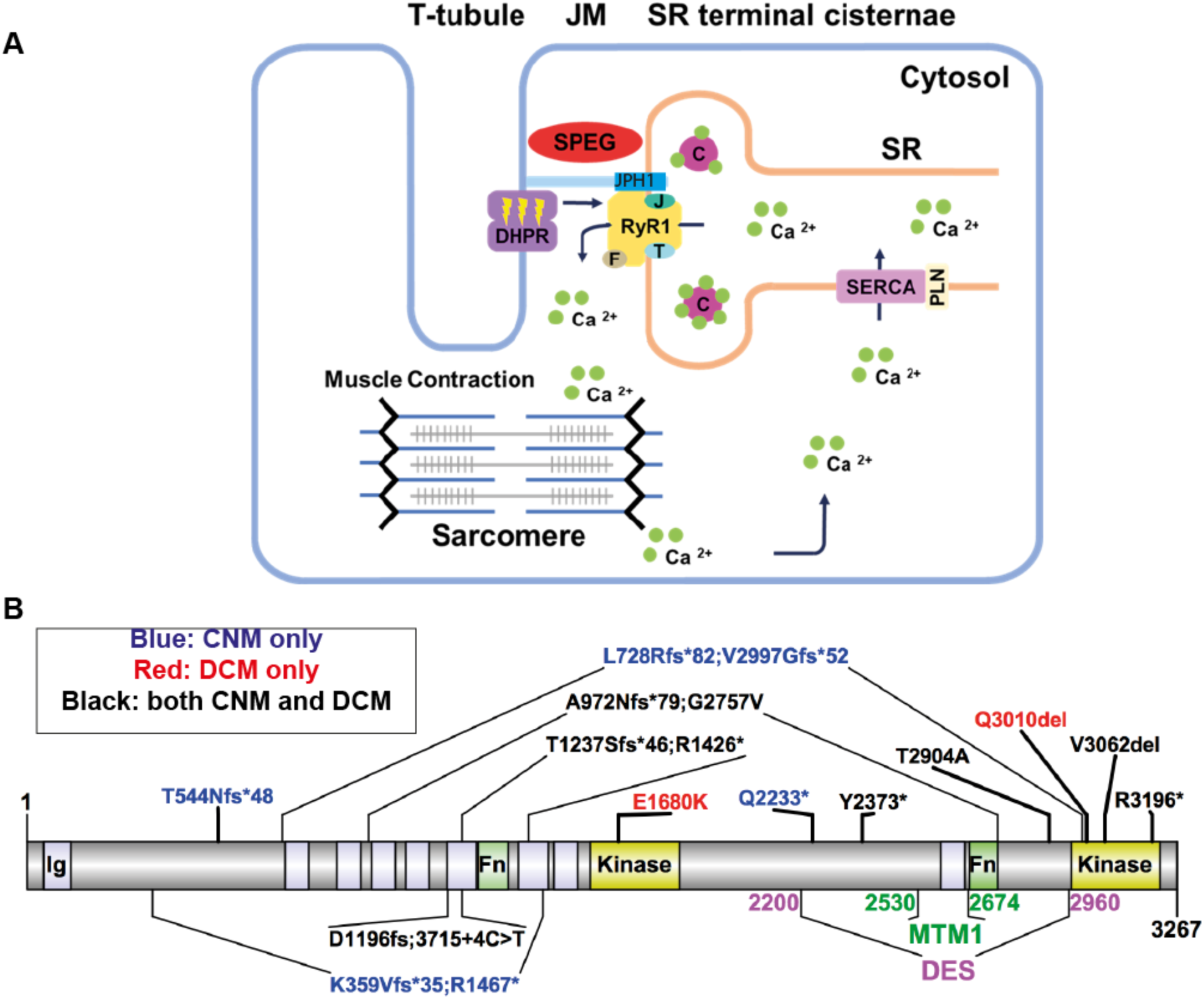
Schematic diagrams of (A) excitation-contraction coupling (ECC) at the triad and (B) *SPEG* domains and pathogenic variants. (**A**) The triads are made of transverse tubules (T-tubules) flanked by terminal sarcoplasmic reticulum (tSR). Junctional membranes (JM) are connected by the interaction of the ryanodine receptor (RyR1) at the tSR, and dihydropyridine receptor (DHPR) at the T-tubules, forming the core components of the ECC machinery. ECC starts when a neuronal action potential arrives via T-tubules and causes a conformational change in DHPR, allowing it to interact with RyR1, causing its activation. As a result, calcium (Ca^2+^) leaves the SR via RyR1 channel opening and moves into the cytosol, promoting sarcomeric contraction. Finally, RyR1 is closed and the calcium transporter SERCA (sarco/endoplasmic reticulum Ca2+-ATPase, regulated by PLN or phospholamban) returns calcium into the SR, where it is largely bound to calsequestrin (C). The terminal SR also contains RyR1 modulators, such as junctophilin-1 (JPH1), FK506-binding protein 1A (F), junctin (J); and triadin (T). SPEG (striated muscle enriched protein kinase) is localized at the triad, but its role remains elusive. (**B**) SPEG contains immunoglobulin domains (Ig, *light purple shaded*), fibronectin domains (Fn, *light green shaded*), and two kinase domains (*yellow shaded*). SPEG directly binds to Myotubularin 1 (MTM1, *dark green*; SPEG 2530∼2674 a.a.) (Agrawal et al., 2014), and Desmin (DES, *magenta*; SPEG 2200∼2960 a.a.) (Luo et al., 2021) at the inter-kinase domain. Pathogenic variants in *SPEG* span different regions of the gene and are typically nonsense mutations that result in decreased SPEG levels. The variants exist either in compound heterozygosity (e.g. K359Vfs*35; R1467*), or homozygosity (e.g. T544Nfs*48). *SPEG* mutations can cause a skeletal muscle disorder only (i.e. centronuclear myopathy or CNM, *blue fonts*), a cardiomyopathy only (e.g. dilated cardiomyopathy, *red fonts*), or both (*black fonts*), with no clear genotype-phenotype correlation. Illustrations were made using Illustrator for Biological Sciences (IBS) (Liu et al., 2015).

*SPEG* encodes a serine/threonine-specific protein kinase that belongs to the Obscurin/MLCK family (Fleming et al., 2021). SPEG contains two serine/threonine kinase domains, two Fibronectin-Type III (Fn) domains, and nine Immunoglobulin (Ig) domains (Fig. 1B). SPEG is predominantly expressed in striated muscles, but also found in the brain (Quick et al., 2017). Previous research suggests that SPEG interacts with proteins involved in the ECC pathway (Agrawal et al., 2014b, Quick et al., 2017, Quan et al., 2019, Huntoon et al., 2018). SPEG is co-localized with RyR1 (Agrawal et al., 2014b), and interacts with MTM1 (Agrawal et al., 2014b) and DES (desmin) (Luo et al., 2020) (Fig. 1B). Bi-allelic variants in *SPEG* have been identified to cause CNM in a small number of families (Agrawal et al., 2014b, Wang et al., 2017, Wang et al., 2018, Lornage et al., 2018, Qualls et al., 2019, Tang et al., 2019, Conlon et al., 2020, Almannai et al., 2021, Zhang et al., 2021) (Fig. 1B). Mutations span different regions of the *SPEG* gene and most are nonsense, resulting in decreased levels of the SPEG protein. Patients with *SPEG* mutations have skeletal muscle weakness as well as dilated cardiomyopathy (DCM). In severe cases, neonatal mortality has been reported (Agrawal et al., 2014b, Liu et al., 2009, Wang et al., 2018, Qualls et al., 2019, Tang et al., 2019). At present, it is unclear how SPEG regulates muscle development, particularly triad formation and/or function, and why mutations in *SPEG* lead to CNM.

SPEG shares high sequence conservation across vertebrates. In humans, a single *SPEG* locus produces a ∼10.8kb transcript that encodes a 3267 amino acid (a.a.) protein (NP_005867.3). In mice, the *Speg* locus includes two alternative transcription start sites that produce four distinct isoforms: *Apeg* (aortic preferentially expressed gene), *Bpeg* (brain preferentially expressed gene), and *Spegα*, and *Spegβ* (Hsieh et al., 2000). Of these, the two longest (*Spegβ*, 3262 a.a., NP_031489.4; and *Spegα,* 2527 a.a., NP_001078839.1) are the striated muscle isoforms, and are well conserved with human SPEG. The striated muscle isoforms are involved in the maturation and differentiation of neonatal cardiomyocytes, while *Spegα* is the predominant isoform involved in skeletal muscle differentiation (Liu et al., 2009, Hsieh et al., 2000). The zebrafish, however, possess two separate *speg* genes, *spega* (Chromosome 6, NP_001007110.1) and *spegb* (Chromosome 9, XP_021334681), each encoding a single Speg transcript. The two zebrafish SPEG proteins (Spega and Spegb) share high sequence conservation with human SPEG, and with both mouse Spegα and Spegβ isoforms.

In this study, we investigated the role of SPEG in skeletal muscle development by generating and characterizing multiple zebrafish *speg* knockout (KO) models (*spega-*KO, s*pegb-*KO, and *spega/spegb* double KO zebrafish). A main advantage of the zebrafish model is that zebrafish larvae can survive up to 7 day-post-fertilization (dpf) without a functional heart (Pelster and Burggren, 1996, Stainier, 2001), allowing for skeletal muscle studies even in the setting of impaired cardiac development. We established zebrafish *speg* KOs using CRISPR/Cas9 directed mutagenesis and confirmed that the resulting mutants exhibited reduced SPEG expression. Phenotypic characterization revealed that *speg*-DKO mutants effectively recapitulate phenotypes reported in *SPEG*-related CNM patients, including abnormal triad structure, disrupted calcium dynamics, and impaired skeletal muscle function. Furthermore, similar to other CNM subtypes, we found that *SPEG* deficiency results in desmin (DES) accumulation and DNM2 upregulation. In total, we established a new zebrafish model suitable for defining the biological role of SPEG in skeletal muscle and for identifying therapies for *SPEG*-related CNM.

## Results

### *spegb* is the primary functional SPEG in zebrafish skeletal muscle

There are two *speg* genes in zebrafish, *spega* and *spegb*. To study tissue expression of *spega* and *spegb* during development, we performed quantitative RT-PCR (RT-qPCR) on total RNAs extracted from the head region (brain, and predominantly cardiac muscle) or tail region (skeletal muscle) of 2 dpf (day-post-fertilization) and 7 dpf zebrafish wild-type (WT) embryos. We detected similar temporal expression pattern of *spega* (Fig. 2A) and *spegb* (Fig. 2B), with significant upregulation observed between 2 dpf to 7 dpf in the head region. We performed whole-mount *in situ* hybridization to compare spatial expression patterns. We observed *spega* expression in the brain and developing neural tube, but not the notochord or somites. In contrast, *spegb* expression was predominantly detected in the chevron-shaped somites (Fig. 2C). These data show that both *spega* and *spegb* are expressed from early development, and that zebrafish skeletal muscle predominantly expresses *spegb*.

**Figure 2.**
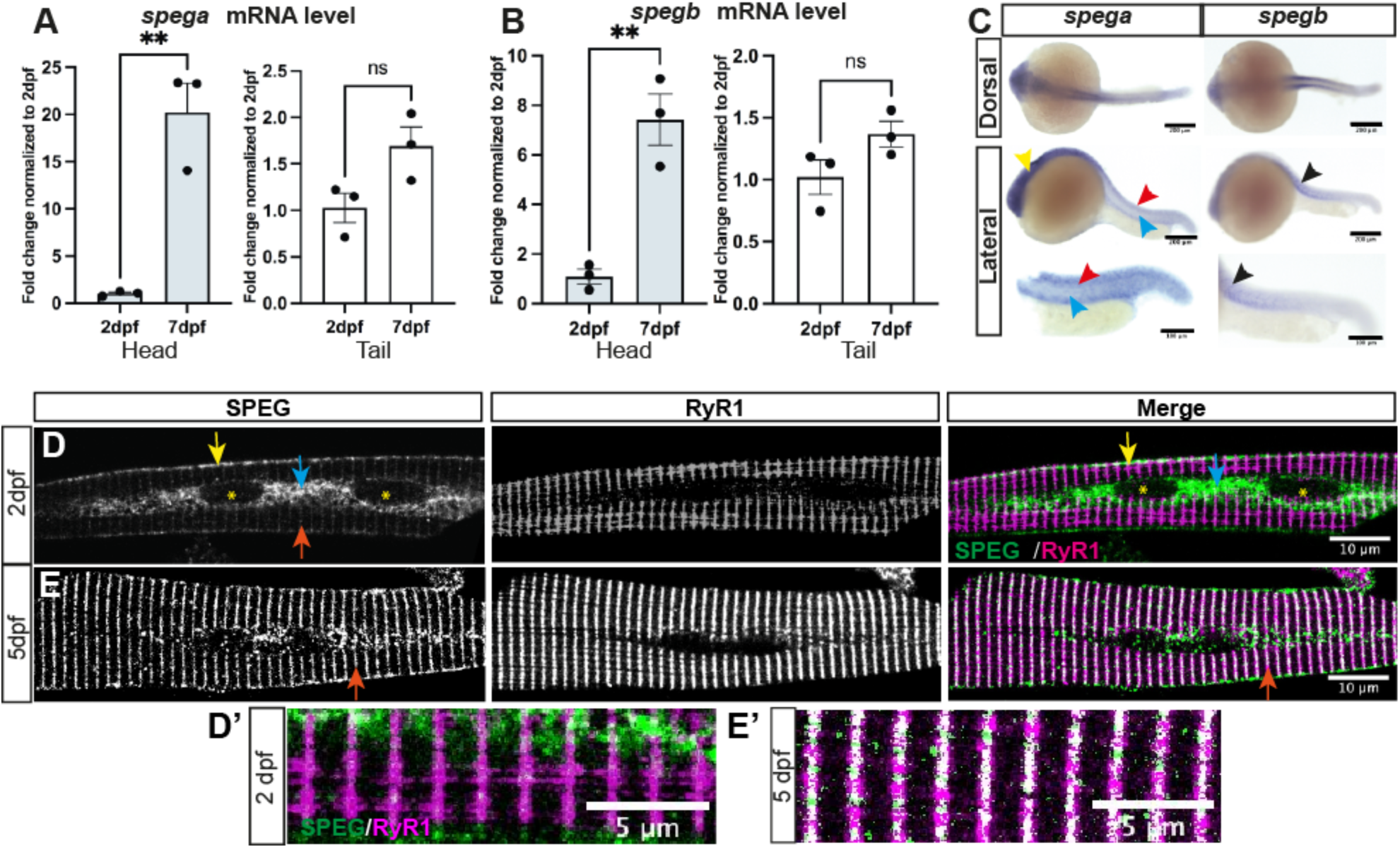
*speg* expression during zebrafish development. There are two SPEG genes in zebrafish, *spega* and *spegb*. Each encodes a single transcript that is highly conserved with human SPEG. (**A-B**) RT-qPCR shows similar temporal expression patterns of *spega* (**A**) vs *spegb* (**B**) from 2 dpf (day-post-fertilization) to 7 dpf. Both SPEG mRNA transcripts are significantly upregulated in the heads (*gray bars*; 20-fold for *spega*, and 7-fold for *spegb*), but stay at relatively similar levels in the skeletal-muscle-predominant tails (*white bars*). Each data point represents the average of technical triplicates, and three independent experiments are included. Columns and error bars represent Mean ± SEM. Two-tailed Student’s *t*-test was performed: **, *P*<0.01; ns, not significant. (**C**) Wholemount *in situ* hybridization using DIG-conjugated RNA probes in 1 dpf embryos show distinct spatial expression patterns of *spega* vs *spegb*. *spega* is predominantly expressed in the developing brain (*yellow arrowhead*) and along the neural tube (*red arrowheads*), but absent from the notochord (*cyan arrowheads*). *spegb* staining, however, is predominantly detected at the chevron-shaped developing somites (*black arrowheads*). In situ images - *scale bars*: 200 μm (upper, middle) or 100 μm (bottom). (**D-E**) Average Z-projections of confocal images showing 2 dpf (**D**) or 5 dpf (**E**) isolated wildtype skeletal myofibres double-stained with anti-RyR1 (34C, DSHB) and anti-SPEG (PA553875, Invitrogen). (**D and D’**) At 2 dpf, SPEG is predominantly localized at the sarcolemma (*yellow arrows*) and perinuclear regions (*cyan arrows*; nucleus: *yellow asterisks*) with weak expression in transverse striations (*orange arrows*), whereas RyR1 is localized in transverse striations labeling the terminal sarcoplasmic reticulum (SR). Little co-localization (*white* in Merge) of SPEG (*green*) and RyR1 (*magenta*) can be observed at this stage. (**E and E’**) At 5 dpf, SPEG expression becomes restricted in the transverse striations (*orange arrows*) that overlap with RyR1. IF images - *scale bars*: 10 μm for D and E, 5 μm for D’ and E’.

### Distinct subcellular localizations of SPEG during early and late muscle development

To characterize the subcellular localization of SPEG in muscle during development, we stained isolated zebrafish myofibres with a SPEG antibody that recognizes both zebrafish Spega and Spegb proteins. We observed by confocal imaging that Speg is localized to sarcolemmal and perinuclear regions at 2 dpf, with little transverse striations or co-localization with the tSR marker RyR1 (Fig. 2D,D’). However, at 5 dpf, Speg loses its early sarcolemmal/perinuclear localization, and instead shows robust transverse co-localization with RyR1 at the triads (Fig. 2E,E’). These results indicate that Speg translocates to the triad junction during muscle development shortly after initial triad formation.

### Generation of *spega* and/or *spegb* CRISPR/Cas9-mediated knockout lines

To study SPEG function during development, we used CRISPR/Cas9 to target the two zebrafish *speg* genes, generating *spega* or *spegb* knockout (KO) lines. We designed guide RNAs (gRNAs) and tested their *in vivo* cutting efficiency using high resolution melting (HRM) analysis (Fig. S1). The most efficient gRNAs were chosen to generate single mutant lines, with double mutant lines later generated by intercrossing F2 single mutants (Fig. 3A). Sanger sequencing confirmed the genotype of one *spega* mutant as carrying a 10-bp deletion in exon 27 (i.e. *spegaΔ10*), and the genotype of one *spegb* mutant as carrying a 17-bp deletion in exon 26 (i.e. *spegbΔ17*), both of which are predicted to introduce premature stop codons in the inter-kinase region (Fig. 3B). To examine the level of *speg* transcripts, we performed RT-qPCR on total RNA extracted from single mutants, double mutants, and their WT siblings. Although no significant transcript reductions were detected in the single mutants (Fig. S2), we observed significantly decreased *spega* (∼50% of WT) and *spegb* (∼40% of WT) in *spegaΔ10;spegbΔ17* double mutants (Fig. 3C). Due to lack of Speg antibodies suitable for western blot analysis in zebrafish, we quantified relative protein levels in skeletal muscle by measuring the intensity (gray values) of immunostaining in isolated myofibres (Fig. 3D). We observed that anti-SPEG staining intensity is significantly reduced by ∼70% in *spegaΔ10;spegbΔ17* myofibres when compared to WT, confirming the double mutant zebrafish to be a Speg-deficient zebrafish line, or *speg*-DKO.

**Figure 3.**
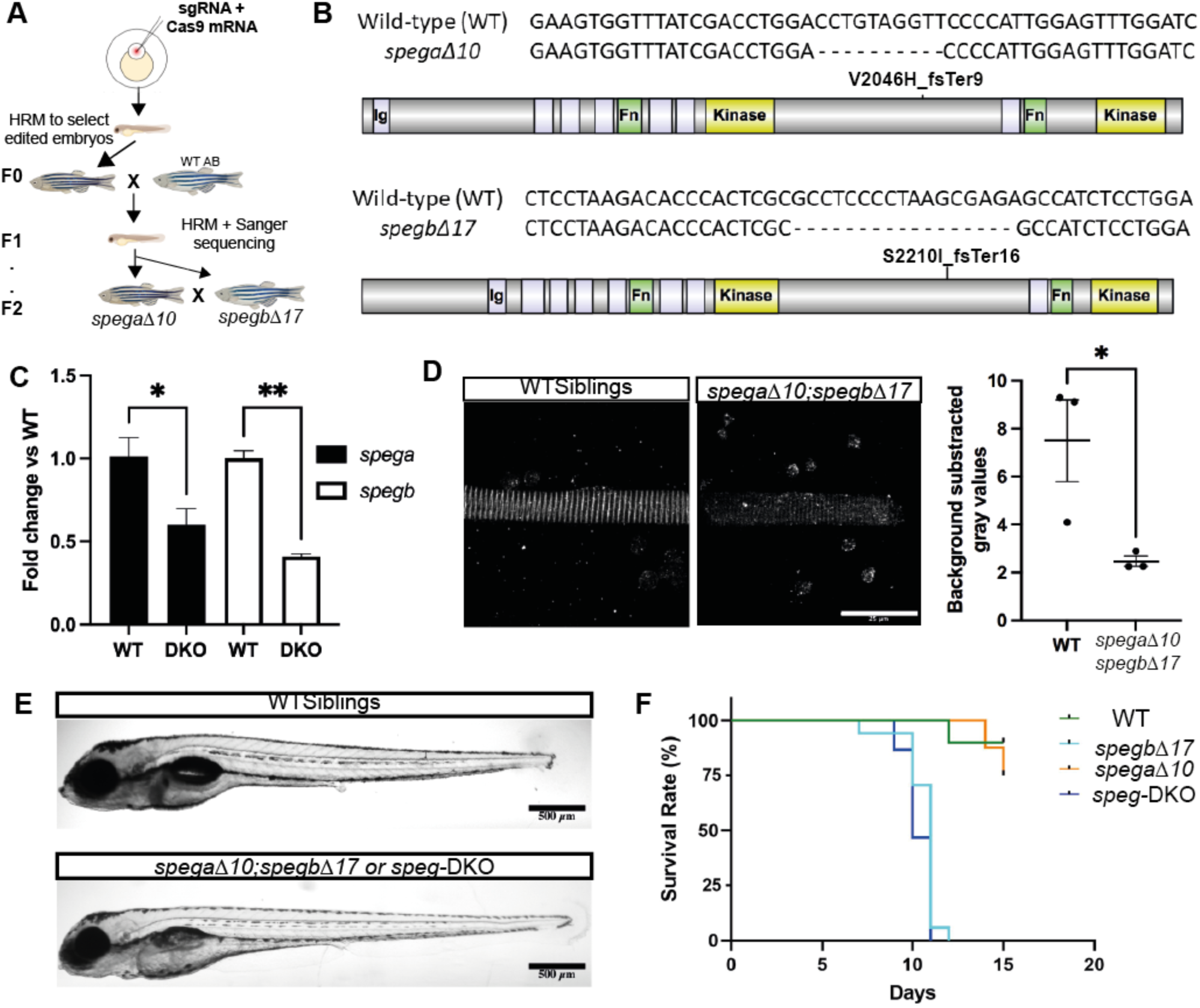
Generation of *speg* knockout zebrafish. (**A**) Schematic diagram showing the workflow of generating *speg* single and double CRISPR-Cas9 mutants. gRNAs (against *spega* or *spegb*) showing high editing efficiency (as determined by HRM) were co-injected with Cas9 mRNA into 1-cell stage embryos (AB strain), and the resulting larvae were raised to adulthood. These F0 zebrafish were then outcrossed to wildtype AB, and resulting embryos screened for germline mutation by HRM and Sanger sequencing. Larvae that showed nucleotide changes leading to premature stop codons were raised to adulthood (F1), i.e. *spega* single KO (*spegaΔ10)*, or *spegb* single KO (*spegbΔ17)*. To dilute off-target effects, F1 adults were outcrossed to AB wildtypes to generate F2 embryos and adults. Double KO lines (e.g. *spegaΔ10;spegbΔ17*) were then generated by crossing *spega* F2 heterozygous (+/-) adults to *spegb* F2 heterozygous (+/-) adults,. (**B**) Genotypes and predicted protein products (if any) of the single mutant lines. *spega*-V2046H_fsTer9: V2046>H, frame shift and terminates in 9 amino acids downstream. *spegb*-S2210I_fsTer16: S2210>I, frame shift and terminates in 16 amino acids downstream. (**C**) RT-qPCR analysis shows significant reductions (around 50%) in both *spega* (*black bars*) and *spegb* (*white bars*) mRNA transcript levels in *spegaΔ10;spegbΔ17*. (**D**) Quantification of Speg protein levels as measured by immunofluorescence (IF, anti-SPEG) intensity in 5 dpf myofibres isolated from WT siblings vs *spegaΔ10;spegbΔ17*. Each dot represents the average of gray values (technical triplicates) measured by the square tool in Fiji ImageJ. Three independent experiments were performed, and error bars are shown as Mean ± SEM. Two-tailed Student’s *t*-test: *, *P*<0.05. **, *P*<0.01. *Scale bar*: 25 μm. (**E**) At 6 dpf, morphology was similar between WT siblings and *spegaΔ10;spegbΔ17* (or *speg*-DKO), though deflated swim bladder was noted in *speg*-DKOs. *Scale bars: 500 μm.* (**F**) Representative Kaplan-Meier curve showing reduced survival in *spegbΔ17* (*spega* +/+; *spegb-*KO, *light blue*) and *speg*-DKO (*spega-*KO*; spegb-*KO, *dark blue*), but not in *spegaΔ10* (*spega-*KO; *spegb* +/+, *orange*) or WT siblings (*spega* +/+; *spegb* +/+, *green*). *spegb*-KO and *speg*-DKO larvae have a median survival of 11 and 10 dpf, respectively; and a maximum survival of 12 and 11 dpf, respectively. N=10 WT, n=17 *spegb*-KO, n=16 *spega*-KO, and n=15 *speg*-DKO. Mantel-Cox test: ****, *P*<0.0001.

### *spegb*-KO and *speg*-DKO zebrafish have significantly reduced survival

We generated single and double knockouts from carrier crosses, and analyzed embryos at the F3 generation and beyond. With the exception of deflated swim bladders in the *speg*-DKO zebrafish, no obvious morphological abnormalities were noted, with embryos and larvae appearing of normal size and without clear bend or curvature to the body shape (Fig. 3E). However, survival was significantly reduced. Using Kaplan-Meier based methodology, we observed that *spegbΔ17* and *spegaΔ10;spegbΔ17* (or *speg*-DKO) died at a median age of 10 dpf (Fig. 3F). On the other hand, *spegaΔ10* survived to adulthood, and could successfully reproduce.

To validate the specificity of the phenotypes that we observe, we generated additional *spega* and *spegb* mutant lines, and examined the survival of *spegaΔ5;spegbΔ8,* another *speg* double mutant line. Specifically, we created lines that induce premature stop codons in *spega* (exon 5, *spegaΔ5*) and *spegb* (exon 8, *spegbΔ8*). These lines are identical in appearance to *spegaΔ10* and *spegbΔ17*, and we observed similar reduced survival rates in *spegbΔ8* and *spegaΔ5;spegbΔ8* (Fig. S3). Similarly reduced survival was seen with *spegbΔ8 +/-;spegbΔ17 +/-* compound heterozygotes (data not shown). These data confirm that the phenotypes we describe are due to *speg* mutation and not to off-target effects. We focused the majority of the remaining analyses on *speg*-DKO zebrafish, and specifically *spegaΔ10;spegbΔ17*, as they best genetically and phenotypically model the human disease.

### *speg*-DKO zebrafish have normal cardiac function

Patients with *SPEG* mutations manifest both skeletal muscle and cardiac (e.g. dilated cardiomyopathy) abnormalities, and heart dysfunction could explain the early lethality seen in our *speg* zebrafish. Therefore, we examined *speg*-DKO embryos and larvae for signs of cardiac abnormalities. One change associated with cardiomyopathy in zebrafish is pericardial edema, caused by disrupted blood flow and decreased contractile function (Huttner et al., 2018). No evidence of pericardial edema was observed in *speg* zebrafish. We also did not detect any obvious alterations in heart rate or cardiac rhythm in *speg*-DKO zebrafish. Therefore, no clear cardiac abnormalities were present in *speg*-DKO zebrafish, suggesting that the early lethality is due to skeletal muscle dysfunction.

### *speg* deficiency results in alterations in T-tubules and triads

To determine if *speg* deficiency leads to skeletal muscle phenotypes that mirror human CNM, we first evaluated the expression pattern of key triad proteins using immunofluorescence (IF) in myofibres isolated from 2 dpf (Fig. 4A-4D) and 5 dpf (Fig. 4A’-4D’) WT vs *speg*-DKOs. In WT myofibres, both RyR1 and DHPR form transverse striations (i.e. triads), as does the sarcomeric Z-disk protein α-actinin, while SERCA1 forms both transverse and longitudinal striations reflective of the SR membrane network. In *speg*-DKO myofibres, however, we observed fragmentation of this striated RyR1 and DHPR pattern, with occasional RyR1 mislocalization to the sarcolemma (Fig. 4A), consistent with abnormal early triad development and/or RyR1/DHPR decoupling. To further dissect this, we performed electron microscopic (EM) analyses and found that triad density (the number of triads per 60 μm^2^) decreased significantly in myofibres from *speg*-DKO zebrafish (Fig. 4F,F’,G), with many triads appearing structurally abnormal (Fig. 4E’). These findings indicate that *speg* deficiency leads to triad loss/abnormality. Interestingly, transverse, but not longitudinal, SERCA staining is depleted in *speg*-DKO myofibres from 5 dpf *speg*-DKO zebrafish (Fig. 4C,C’), while α-actinin staining remained normally striated (Fig. 4D,D’), consistent with the normal overall sarcomeric organization observed under EM (Fig. 4E,F). Together, these results indicate that SPEG is essential for triad formation/stability in skeletal muscle.

**Figure 4.**
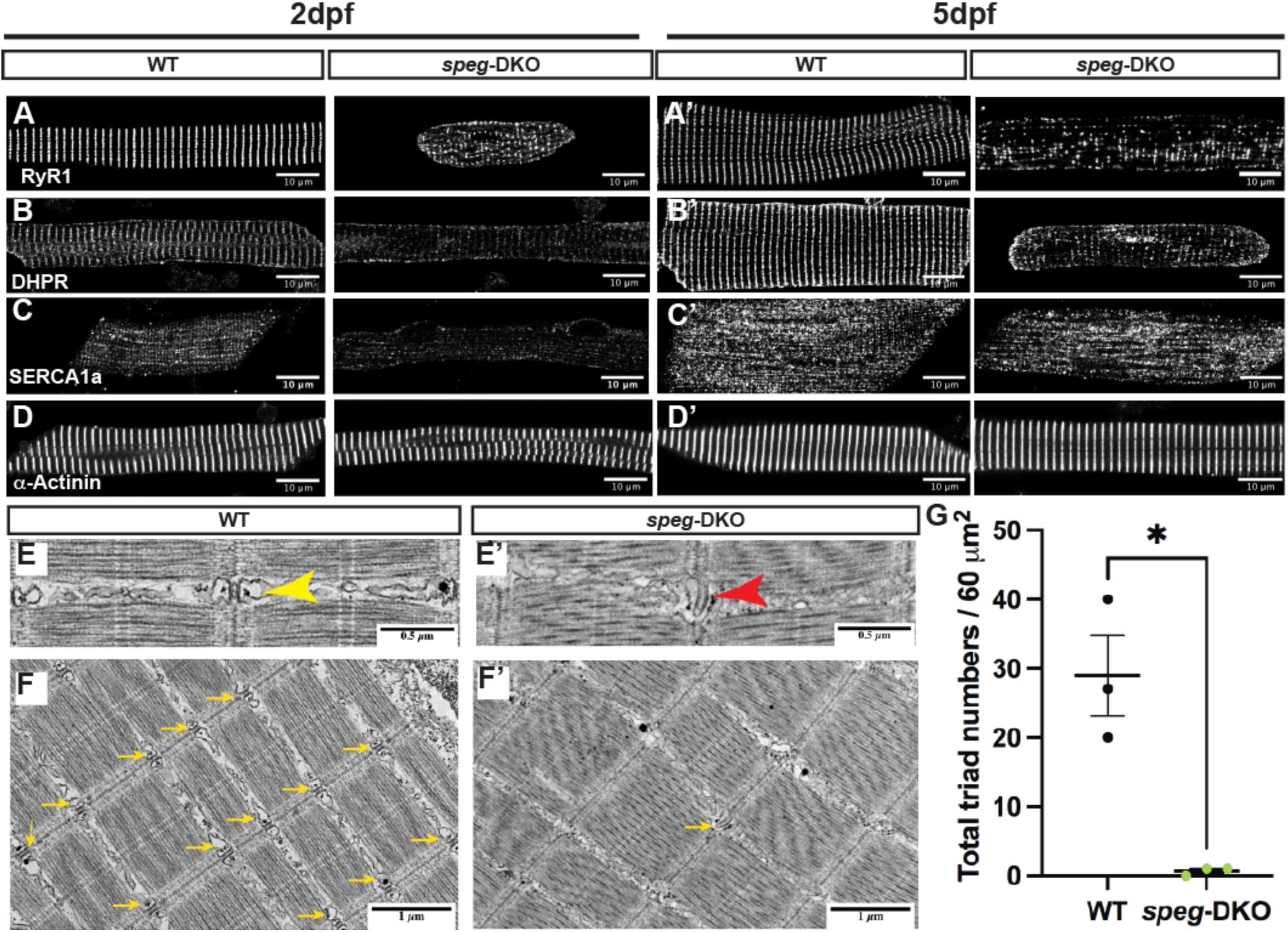
*speg* deficiency in zebrafish disrupts triad protein organization and triad ultrastructure, leading to reduced triad numbers. Immunofluorescence staining was performed on 2 dpf (**A-D**) and 5 dpf (**A’-D’**) isolated myofibres. Average projections of confocal Z-stacks show disrupted transverse pattern of RyR1 (**A** and **A’**), DHPR (**B** and **B’**), and SERCA1a (**D** ad **D’**) in *speg*-DKO starting from 2 dpf, while α-Actinin (**D** and **D’**) is not affected. IF images - *scale bars*: 50 μm. (**E-F**) Electron micrographs of 7 dpf WT and *speg*-DKO muscles. (**E**) In WT zebrafish skeletal muscle, normal triads are physically above the sarcomeric Z-disks, and composed of centrally-located T-tubules flanked by terminal sarcoplasmic reticulum (*yellow arrows*). (**E’**) Triads in *speg*-DKO appear structurally disrupted, losing the obvious tSR/T-tubule/tSR pattern (*red arrows*). More importantly, (**F**) the majority of sarcomeric Z-disks in *speg*-DKO do not have adjacent triads (*yellow arrows*). EM images – *scale bars:* (E) – 0.5 μm; (F) – 1 μm. (**G**) The total number of triads per 60 μm^2^ (under electron microscopy) was significantly reduced in *speg*-DKO. Each dot represents the average of technical triplicates, and three biological replicates are included. Error bars are shown as Mean ± SEM. Two-tailed Student’s *t*-test: *, *P*<0.05.

### *speg* deficiency disrupts ECC and impairs zebrafish swimming performance

We then set out to examine the effects of *speg*-DKO on excitation-contraction coupling (ECC) and RyR1 function. ECC was quantified in single myofibres by measuring intracellular calcium dynamics during electrical stimulation and ligand-induced RyR1 Ca^2+^ release was assessed following local caffeine application. In brief, single myofibres were isolated from genotyped larvae at 7 dpf, loaded with Ca^2+^ sensitive dye (fluo-4AM) (Fig. 5A), and then exposed to a series of electric stimulations (1 Hz, Fig. 5B; and 10 Hz, Fig. 5C,D) followed by 30-second application of 10 mM caffeine (Fig. 5E). Peak changes of fluo-4 fluorescence (myoplasmic free Ca^2+^) were recorded and normalized to background (ΔRatio). Both myofibres from both *spegb*-KO (bKO) and *spega/b*-DKO (DKO) zebrafish exhibited significantly reduced electrically-evoked and caffeine-induced Ca^2+^ release (∼50% reduction in ΔRatio) compared to that observed for myofibres from WT zebrafish (Fig. 5B-5E). These results indicate that *speg* deficiency results in reduced ECC and RyR1 function in skeletal muscle, consistent with the observed alterations in T-tubules and triads. To examine whether *speg*-DKO alters motor performance, we conducted whole zebrafish swimming assays using ZebraBox (Viewpoint, France) (Fig. 5F). As early as 3 dpf, *speg*-DKO zebrafish traveled significantly less distance (29-49% reduction) compared to their WT siblings (Fig. 5G). By 7 dpf, *speg*-DKO zebrafish have a complete absence of movement. Of note, *spegb* mutants also demonstrated abnormal swim behavior, consistent with *spegb* representing the primary functionally relevant *SPEG* paralog in skeletal muscle.

**Figure 5.**
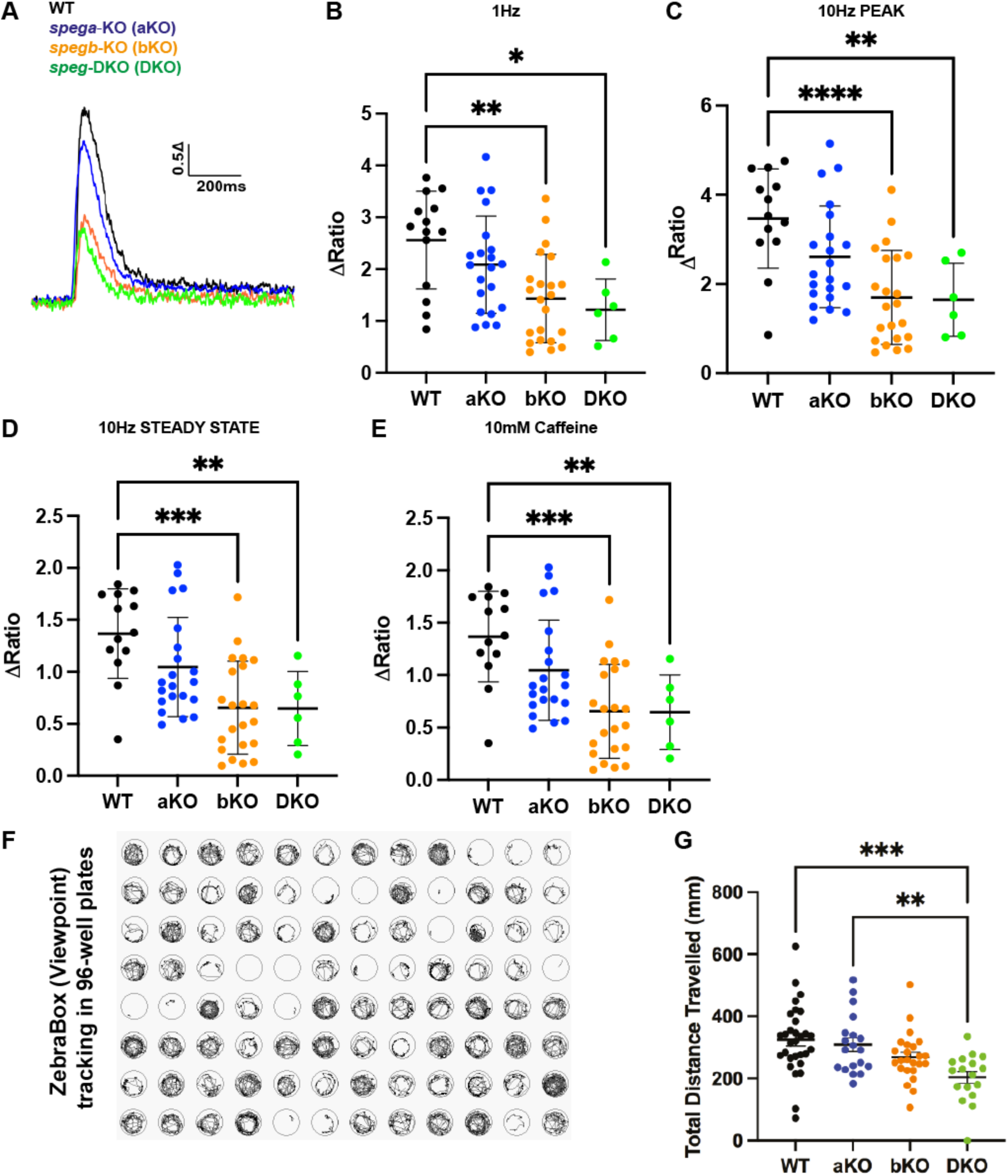
*speg* deficiency in zebrafish disrupts excitation-contraction coupling (ECC) in isolated myofibres and impairs overall muscle performance. (**A-E**) Cytosolic Ca^2+^ transients in 7 dpf isolated myofibres were measured using a Ca^2+^ sensitive dye fluo4-AM: (**A**) A representative diagram of Ca^2+^ transient traces. (**B**) Five-twitch 1 Hz stimulations were followed by (**C-D**) one 10 Hz stimulation, and (**E**) 30 sec of 10 mM caffeine. Overall, *spegb*-KO (bKO) and *speg*-DKO (DKO) showed significantly reduced ECC compared to WT and *spega*-KO (aKO). (**F**) A representative image of swim trace tracking using ZebraBox in a 96-well plate setting, by which total distance travelled (mm) was quantified for each 3 dpf zebrafish per well. (**G**) Both *spegb*-KO and *speg*-DKO swam significantly less distance than WT and *spega*-KO. All statistical analyses include three independent experiments: for fluo4-AM experiments (B-E), each dot represents one myofibre, at least n=5 myofibres per group per experiment; for swimming assay (G), each dot represents a 3 dpf zebrafish, at least n=5 zebrafish per group per experiment. Error bars are shown as Mean ± SEM. One-way ANOVA: *, *P*<0.05; **, *P*<0.01; ***, *P*<0.001; ****, *P*<0.0001. Note – WT (black line or dots); aKO: *spega*-KO (blue line or dots); bKO: *spegb*-KO (orange line or dots); DKO – *spega/b* double KO (green line or dots).

### *speg*-DKO shows abnormal Desmin accumulation and upregulation as early as 5 dpf

Previous work identified DES (desmin) (Luo et al., 2020) and myotubularin (MTM1) (Agrawal et al., 2014a) as SPEG binding partners in skeletal muscle, and aberrant DES aggregation in *Mtm1* KO mice (Hnia et al., 2011a, Luo et al., 2020). We thus compared DES expression in a series of CNM zebrafish models at stages where muscle phenotypes are fully penetrant: 1) *speg*-DKO at 5-7 dpf (Figs. 3-5), 2) *mtm1*-KO at 7 dpf (Sabha et al., 2016), and 3) *DNM2*-S619L-eGFP transgenics at 3 dpf (Zhao et al., 2019). To examine DES localization, we stained isolated myofibres with a DES antibody that picks up both Desmin-a and Desmin-b in zebrafish (Kayman Kürekçi et al., 2021). In myofibres from WT zebrafish, DES forms transverse striations marking the sarcomeric Z-disks, and is also localized around the nucleus and below the sarcolemma (Fig. 6A,D). In both *speg*-DKO (Fig. 6B) and *DNM2*-S619L (Fig. 6E) this clear striated pattern is lost, and abnormal accumulation is observed longitudinally within the middle of the myofibre and below the sarcolemma. In *mtm1*-KO, mild accumulation of DES is observed longitudinally within the middle of the myofibre without an obvious loss of transverse striations (Fig. 6C).

**Figure 6.**
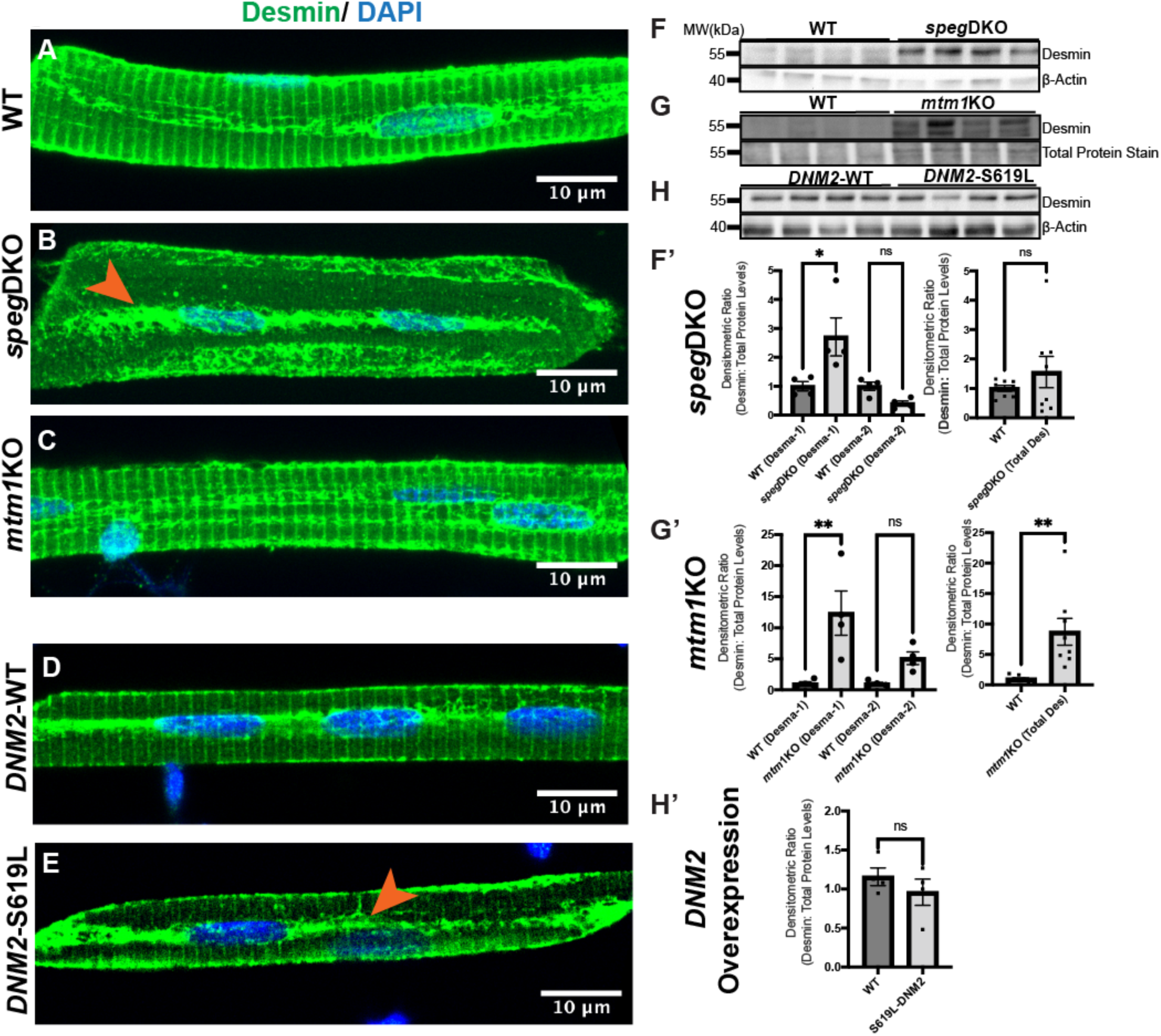
Desmin subcellular localization and protein levels in zebrafish models of CNM. **(A-D)** Myofibres were isolated at 5 dpf (*speg*-DKO), 7 dpf (*mtm1*-KO) and 3 dpf (*DNM2* overexpression), and stained with anti-Desmin (*green*; D8281, Sigma) and DAPI (*blue,* nucleus). (**A**) At 5 dpf, Desmin is normally localized to the sarcolemma, perinucleus, and the sarcomeric Z-disks (as transverse striations). (**B**) In *speg*-DKO (5 dpf), Desmin is predominantly localized to the perinucleus (*orange arrowhead*). (**C**) In *mtm1*-KO (7 dpf), Desmin localization appears similar to wildtype (WT) siblings. (**D**) WT-*DNM2*-EGFP overexpressing myofibres show similar Desmin staining pattern to non-transgenic WTs, while (**E**) S619L-*DNM2*-EGFP overexpressing myofibres show loss of Desmin in transverse striations, with Desmin localization predominantly at the perinucleus (*orange arrowhead*). IF images - *scale bars*: 10 μm. (**F-H, F’-H’**) Western blot analysis using whole zebrafish lysates shows Desmin upregulation in (**F** and **F’**) 5 dpf *speg*-DKO (by 2-3 fold) and in (**G** and **G’**) 7 dpf *mtm1*-KO (by 5-10 fold) when compared to WT siblings, but not in (**H** and **H’**) 3 dpf *DNM2*-S619L zebrafish when compared to *DNM2*-WT controls. Each lane (**F-H**) or each dot (**F’-H’**) represents n=25 zebrafish (40 μg of total proteins), four lanes represent four independent experiments. Densitometry was measured using Fiji ImageJ. Desmin protein levels were normalized to β-actin loading controls in *speg*-DKO and *DNM2*, or to REVERT total protein stains in *mtm1*-KO (as β-actin level is changed by the lack of MTM1). Error bars are shown as Mean ± SEM. Two-tailed Student’s *t*-test: *, *P*<0.05; **, *P*<0.01; ns, not significant.

To determine if overall DES expression levels are altered, we performed western blot analysis on whole zebrafish protein lysates. In WT sibling controls (Fig. 6F, 5 dpf; and Fig. 6G, 7 dpf), we observed two DES bands (an upper band ∼55 kDa and a lower band ∼50 kDa), consistent with a previous report in adult zebrafish muscle (Kayman Kürekçi et al., 2021). In the mutants, we detected significant DES upregulation in both 5 dpf *speg*-DKO zebrafish (2-2.5 fold, Fig. 6F’) and 7 dpf *mtm1*-KO zebrafish (9-12 fold, Fig. 6G’) compared to WT controls. Conversely, no difference in DES expression was detected between 3 dpf *DNM2*-WT and *DNM2*-S619L zebrafish (Fig. 6H,H’). Of note, only a single DES band at ∼55kDa was observed in the *DNM2* experiments, most likely due to the earlier developmental time point (3 dpf) used in these studies. In summary, increased longitudinal DES accumulation was observed across all three models, while increased DES expression was observed only with 5 dpf *speg*-DKO and 7 dpf *mtm1*-KO zebrafish.

### Dnm2 is upregulated and disorganized in skeletal muscle of *speg*-DKO zebrafish

Upregulation of endogenous DNM2 protein is a common phenotype observed in non-*DNM2*-related CNM models including mice lacking *Mtm1* (Cowling et al., 2014b) and *Bin1* (Cowling et al., 2017b). Thus, we evaluated Dnm2 protein expression in our *speg*-DKO zebrafish and *mtm1*-KO zebrafish. We first studied Dnm2 localization by performing IF on isolated myofibres using an antibody specifically against zebrafish Dnm2. In WT myofibres (5 dpf), Dnm2 staining appears in transverse striations (Fig. 7A). In *speg*-DKO (Fig. 7B) and *mtm1*-KO (Fig. 7C), while Dnm2 staining forms similar striated patterns, occasional Dnm2 aggregates can be observed along the striations. However, these patterns are distinct in appearance and localization from the large sarcolemmal DNM2-EGFP aggregates previously observed with *DNM2*-S619L overexpression (Zhao et al., 2019). To examine Dnm2 protein levels, we performed western blot analysis in 5 dpf *speg*-DKO (Fig. 7D) and 7 dpf *mtm1*-KO (Fig. 7E) zebrafish using whole zebrafish protein lysates. We detected two Dnm2 bands at ∼100 kDa in WT siblings, with a significant (∼2-3 fold) increase in the upper band in both *speg*-DKO (Fig. 7D,D’) and *mtm1*-KO zebrafish (Fig. 7E,E’), consistent with DNM2 upregulation in *SPEG*- and *MTM1*-CNM. Together, our results demonstrate that DNM2 is upregulated in *SPEG*- and *MTM1*-CNM in the absence of the detectable sarcolemmal DNM2 aggregation seen with *DNM2*-CNM.

**Figure 7.**
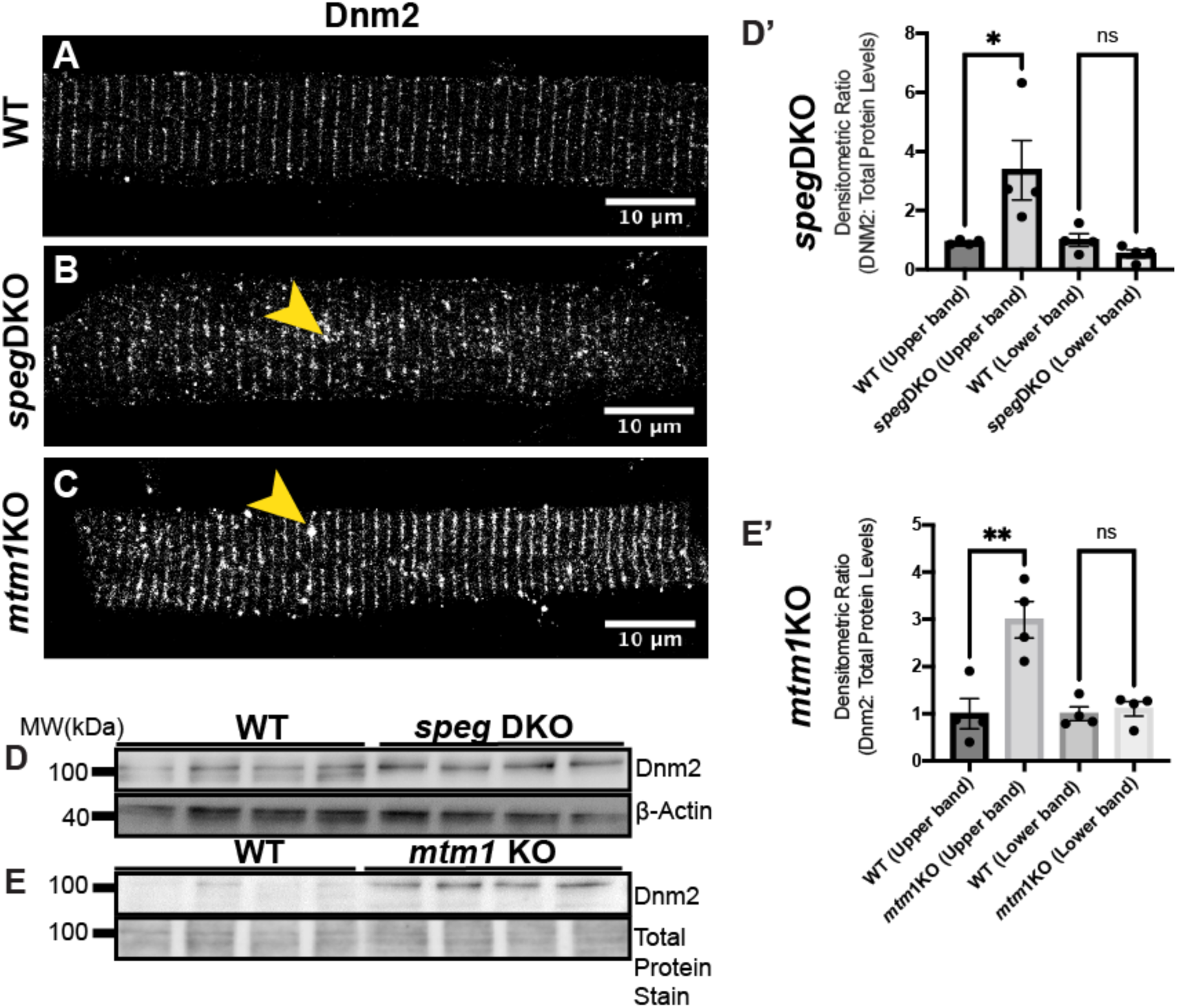
Dnm2 protein is upregulated in both *speg*-DKO and *mtm1*-KO zebrafish. **(A-C)** Myofibres were isolated at 5 dpf (*speg*-DKO) and 7 dpf (*mtm1*-KO), and stained with anti-Dnm2 (GTX127330, GeneTex). (**A**) In wildtype myofibres (5 dpf), Dnm2 is localized to the triads (transverse striations). Similar striated patterns can be observed for Dnm2 in (**B**) *speg*-DKO and (**C**) *mtm1*-KO, with occasional Dnm2 aggregations observed along the striations (*yellow arrowheads*). IF images – *scale bars*: 10 μm. (**D-E**) Western blot analysis shows increased Dnm2 protein levels in (**D and D’**) 5 dpf *speg*-DKO and (**E and E’**) 7 dpf *mtm1*-KO when compared to WT. Each lane (**D-E**) or each dot (**D’-E’**) represents n=25 zebrafish (40 μg of total protein). Densitometry was accomplished using Fiji ImageJ. Dnm2 protein level was normalized to β-actin loading controls in *speg*-DKO, or to REVERT total protein stains in *mtm1*-KO (as β-actin level is changed by the lack of MTM1). Students *t*-test: *, *P*<0.05; **, *P*<0.01; ns, not significant.

## Discussion

*SPEG* is a novel CNM-causing gene and the only known kinase associated with CNM. The function of SPEG in skeletal muscle remains unclear. In this study, we provide the first characterization of SPEG in zebrafish. We define the developmental expression of *spega/b* mRNA, as well as the subcellular localization of SPEG protein in skeletal muscle fibres. We generated single and double knockouts of zebrafish *speg* paralogs, and show that the double knockout zebrafish exhibit molecular, pathologic, and behavioral phenotypes consistent with the human disease. We also examined potential *SPEG*-CNM pathomechanisms and therapeutic targets, and found that, as with other forms of CNM, *SPEG* deficiency is associated with both DES (desmin) accumulation and DNM2 (dynamin 2) upregulation. In all, we establish a new model of *SPEG*-deficient CNM that will be ideal for studying disease mechanisms and identifying therapies.

There are two *speg* genes in zebrafish, *spega* and *spegb*. Although both share high sequence conservation with human *SPEG* and are expressed from early zebrafish development, our *in situ* hybridization data demonstrate that the zebrafish skeletal muscles predominantly express *spegb*. This is further supported by our *loss-of-function* data, where *spegb* single KOs showed similar triad and calcium transient defects as *spega/b* double knockouts, and also had reduced survival. However, we only detected significantly reduced swimming distance in *spega/b* double knockouts, suggesting that the lack of both paralogs are required to induce the full muscle phenotype in zebrafish that mirrors that of *SPEG*-CNM patients.

Insights into the roles of SPEG during muscle development can be inferred from known SPEG interacting partners identified by yeast two-hybrid and validated in subsequent co-immunoprecipitation experiments, such as Desmin (Luo et al., 2020), and MTM1 (Agrawal et al., 2014b). Desmin is a major intermediate filament protein in muscle that promotes muscle integrity by forming dynamic filamentous networks crosslinking the sarcolemma, sarcomeres, and mitochondria (Paulin and Li, 2004). Emerging evidence has shown that Desmin (DES) is an important CNM disease marker. Abnormal DES accumulation is observed in muscle biopsies from *MTM1*-CNM and *DNM2*-CNM patients (Romero and Bitoun, 2011). Speg-deficient mice (Luo et al., 2020), Mtm1-deficient mice (Hnia et al., 2011b), as well as our *speg*-DKO, *mtm1*-KO, and *DNM*2-S619L zebrafish, all show abnormal DES accumulation and/or overall DES upregulation. Although it remains unknown the cause for DES accumulation, reduced/altered DES phosphorylation disrupts intermediate filament disassembly and promote DES accumulation (Winter et al., 2013). Thus it is tempting to speculate that SPEG regulates DES phosphorylation, and that lack of SPEG leads to DES accumulation because of reduced phosphorylation. Future experimentation will be required to test this potential link, and more generally to define substrates of SPEG’s kinase activity.

The biological consequences of DES accumulation in *SPE*G-CNM remain unclear. Notably, DES accumulation has been associated with mitochondrial defects in *Mtm1* mouse muscles (Hnia et al., 2011b), and mitochondrial defects along with triad disruption are a key aspect of the CNM disease process (Zanoteli et al., 2009, Lawlor et al., 2016, Muñoz et al., 2020a). It will thus be of interest in the future to examine mitochondrial dynamics in SPEG knockouts. Notably, DES accumulation is also present in desminopathies, a subset of myofibrillar myopathies caused by autosomal dominant *DES* mutations, which have neither triad defects nor other histopathologic features of CNM, indicating that DES accumulation is unlikely a direct cause for the triad defects observed in CNM. Furthermore, in contrast to desminopathies, we did not observe myofibril dissolution (as indicated by normal α-actinin striations) in our CNM zebrafish models, further supporting distinct disease pathways of the two myopathies, consistent with the clinical and histopathological differences between CNM and desminopathy patients.

The triad defects observed in *SPEG*-CNM are potentially due to disrupted SPEG/MTM1 interaction. *MTM1, DNM2*, and *BIN1* all encode components of the endocytic machinery. Since mutations in each of these genes cause CNM with defective triads, membrane trafficking is considered a key aspect of triadogenesis (Dowling et al., 2008). One theory on triad biogenesis is that T-tubule invagination is first initiated by membrane deformation promoted by BIN1 and then completed by DNM2-mediated (Nicot et al., 2007b) membrane fission (Picas et al., 2014, Cowling et al., 2017b). However, the precise roles of SPEG and MTM1 (a phosphatase that acts to dephosphorylate 3-position phosphoinositides (PI[3]P and PI[3,5]P2)) in this pathway are unclear. In *Mtm1*-KO and *Bin1*-KO mice, the level of Dnm2 protein is upregulated (Cowling et al., 2014a, Cowling et al., 2017a). In our *mtm1*-KO and *speg*-DKO zebrafish, we observed similar increases in Dnm2 protein expression, demonstrating that DNM2 is also upregulated in *SPEG*-CNM. SPEG/MTM1 interaction therefore may regulate triad biogenesis by regulating DNM2 levels and/or activity. The mechanisms leading to DNM2 upregulation in *SPEG*-CNM (this study), *BIN1*- (Cowling et al., 2017a), and *MTM1*-CNM (Cowling et al., 2014a) remain unclear, although lack of BIN1 inhibition (Cowling et al., 2017a), dysregulated microRNA (Liu et al., 2011, Chen et al., 2020), and altered post-translational modifications (e.g. SPEG phosphorylation) (Kusić et al., 2020) have been proposed. Moreover, abnormal DNM2 aggregates are observed (Chin et al., 2015, Zhao et al., 2019) or suggested (Muñoz et al., 2020b) in *DNM2*-CNM caused by hyperactive *DNM2* mutations. However, unlike *DNM2*-S619L zebrafish (which model *DNM2*-CNM, we did not observe sarcolemmal Dnm2 aggregates in myofibres from *speg*-DKO or *mtm1*-KO zebrafish, indicating a level of variability in DNM2 abnormalities across different CNM subtypes. Interestingly, DNM2 reduction has been used as an effective strategy in treating muscle phenotypes of *MTM1*- and *BIN1*-CNM mouse models (Tasfaout et al., 2018). It will thus be important to test reducing DNM2 levels (e.g. via genetic ablation, anti-sense oligonucleotides, or manipulating upstream interactions/events) in models of *SPEG*-CNM.

Given that the subcellular localization of SPEG changes rapidly from being perinuclear/sarcolemmal at 2 dpf to located within the triad junction at 5 dpf, SPEG may play distinct roles in developing vs mature muscles. The co-localization of SPEG/RyR1 in mature myofibres suggests that SPEG and RyR1 may interact (similar to SPEG/RyR2 interaction in cardiac muscles) (Quick et al., 2017), such that SPEG promotes phosphorylation events that modulate RyR1 channel function/stability (Jungbluth et al., 2018, Witherspoon and Meilleur, 2016). Of note, it remains unclear how SPEG is recruited to each cellular compartment. Some of these questions could be addressed in future studies that compare RyR1 phosphorylation/interactomes in developing vs mature muscles from WT and *speg*-DKO zebrafish. Lastly, in the absence of any effective treatments for *SPEG*-CNM, our *speg*-DKO zebrafish will enable large-scale chemical screens to identify novel small-molecules/pathways to treat the devastating disease.

## Conclusions

In conclusion, we describe the generation and characterization of the first zebrafish model of *SPEG*-CNM. We demonstrate that *speg*-DKO zebrafish faithfully recapitulate multiple features of *SPEG*-CNM, and identify changes in DES and DNM2 expression/localization as critical (and conserved) disease markers that can be dissected in future studies.

## Acknowledgements

The authors gratefully thank Dr. Yukari Endo for reviewing the manuscript, and Dr. Ramil R Noche for useful discussions and insightful suggestions; Alejandro Salazar, and Elyjah Schimmens for zebrafish maintenance (SickKids Zebrafish Facility); SickKids Imaging facility and SickKids Nanoscale Biomedical Imaging Facility for helps with microscopic training and analysis.

## Author contributions statement

*Conceptualization* – KGE, ICS, and JJD. *Methodology* – KGE, SG, LG, JV, RTD, MZ, and JJD. *Validation* – KGE, SG, LG, MZ, and JJD. *Formal analysis* – KGE, SG, LG, ICS, RTD, MZ, and JJD. *Investigation* – KGE, SG, LG, JV, and MZ. *Visualization* – KGE, SG, and MZ. *Resources* – RTD and JJD. *Writing (Original Draft Preparation)* – KGE. *Writing (Review & Editing)* – KGE, SG, LG, ICS, RTD, MZ, and JJD. *Supervision* – ICS, RTD, MZ, and JJD. *Project administration* – MZ and JJD. *Funding acquisition* – ICS, RTD, and JJD.

## Materials and Methods

### Zebrafish maintenance

Zebrafish (AB strain) were raised and maintained at 28.5°C at the Zebrafish Facility at the Hospital for Sick Children, Toronto, ON, Canada. Experiments were performed on zebrafish embryos and larvae from the one-cell stage up to 15 dpf. All zebrafish procedures were performed in strict accordance with the Animals for Research Act of Ontario and the Guidelines of the Canadian Council on Animal Care.

### Quantitative Real-Time PCR (RT-qPCR)

Total mRNA was isolated from 2 dpf or 7 dpf zebrafish homogenates using the RNeasy Mini Kit (Qiagen, Cat# 74104) and reverse-transcribed with SuperScript VILO (ThermoFisher, Cat# 11755050). Approximately, 10-25 embryos were collected for each condition, in triplicates. Head and trunk dissections were done to cut at the region indicated at the diagram (Fig. S4). RT-qPCR was performed using Platinum SYBR Green reagent (ThermoFisher, Cat# 11744500) and the Step-One-Plus Real-Time PCR System (Applied Biosystems). All reactions were performed in technical triplicates and the results represent biological triplicates. The zebrafish peptidylprolyl isomerase B gene, *ppib*, was used as the endogenous control. Primers used are as follows: *spega* forward 5’-ACAAAGAGATTGGCAGAGGGG-3’, reverse 5’-ACTCTCGCAATGCACAAGTC-3’; *spegb* forward 5’-CAACAACAAGTACGGCAGCG-3’, reverse 5’-TGCAAATCGAGGAGTCTCGC-3’; and *ppib* forward 5’-ACCCAAAGTCACGGCTAAGG-3’, reverse 5’-CTGTGGTTTTAGGCACGGTC-3’. Protocol Conditions for RT-qPCR are the following: 95°C 10’ → (95°C 15” → 60°C 1’) (x40 cycles) → [95°C 15” → 60°C 1 ’ → 95°C 15”] [Melt Curve].

### Wholemount *in-situ* hybridization (ISH)

Probes were generated via PCR amplification from 2 dpf total cDNA synthesis using SuperScript VILO (ThermoFisher, Cat# 11755050). Primers used were as follows: *spega* forward 5’-AAGAAGCAAGCTCACCCACA-3’, reverse 5’-AAGTCAAGGTCTGTCGACGC-3’; *spegb* forward 5’-CGAAACTCACACGGGGAAGA-3’, reverse 5’-GACTGTGATGCTCAAGGGCT-3’. DIG-labeled *in-situ* probes were synthesized using DIG RNA Labeling kits (Roche, Cat# 11277073910). RNA *in-situ* hybridization was carried out as previously described (Thisse and Thisse, 2007). Briefly, embryos were fixed in 4% paraformaldehyde (PFA) and then dehydrated in 100% Methanol. Embryos were then permeabilized and incubated with digoxigenin-labeled antisense RNA probes at a final concentration of 300 ng / 200 μL in hybridization solution. Hybridizations of the probe with the RNA were detected with an alkaline phosphatase-conjugated antibody (Anti-Digoxigenin-AP, Fab Fragments, 1:5000, Roche, Cat# 11093274910). Finally, stained embryos were cleared overnight in a 70% glycerol solution with 30% PBSTw (0.1% Tween 20 in PBS), and imaged under a Leica M205FA stereomicroscope.

### Generation of zebrafish *speg* mutants

The program Chopchop (http://chopchop.cbu.uib.no/) (Montague et al., 2014) was used to design each of the guide RNAs (gRNAs) used in this project. According to Chopchop, there were no predicted off-targets for the gRNAs tested. Next, 50 to 100 one-cell stage WT embryos were injected with the gRNA (150pg per embryo) and Cas9 mRNA (100pg per embryo) with a Picopump (World Precision Instruments). Strong gRNAs were identified by isolating genomic DNA from 24 individual injected embryos and 3 uninjected embryos at 3-5 dpf. DNA was digested with 1 µg/µl Proteinase K and all embryos were genotyped using HRM analysis and Sanger Sequencing. HRM analysis was performed on a Roche Lightcycler 96. Once we identified gRNAs that were cutting at the desired genomic region, potential founders (F0) were outcrossed to WT AB zebrafish. In-cross progeny from the F3 and F4 generations were used for the characterization of the *speg* mutant phenotype. The targets in this study were: *spega* 5’-CTATCGACCTGGACCTGTAGG-3’ at exon 27, and *spegb* 5’-ATGGCTCTCGCTTAGGGGAGG-3’ at exon 26. Primers used for HRM are as follows: *spegaΔ10* at exon 27 forward 5’-AAGAAAGCTCACCGGTTCCC-3’, reverse 5’-TGGACAGACTTGGATTTTTCCT-3’; and *spegbΔ17* at exon 26 forward 5’-CCCTCCCAAAGAGCCAAGTC-3’, reverse 5’-GCGAGCTTCAAAAACCTCCT-3’. Primers used for PCR prior to Sanger sequencing are as follows: *spegaΔ10* forward 5’-ACACCATTACCGACACCAGTT-3’, reverse 5’-TACGTCGCACTGCAAGGAC-3’; and *spegbΔ17* forward 5’-TCTTTTCTCGGGTTGCCTCC-3’, reverse 5’-GAGAGTCGCCTCATGAACCC-3’.

### Morphological assessment and survival analysis

Fish were examined daily for morphological abnormalities, and imaged at 6 dpf using Zeiss Axio Zoom V16 microscope (16x, 10 ms exposure). Survival analysis was performed using Kaplan-Meier methods. Briefy, 3-5 dpf larvae were fin clipped and then grouped by genotype after HRM analysis. Survival count and health check was performed daily until they reached adulthood. The number of live embryos was plotted across time and analyzed using GraphPad Prism version 8 (GraphPad Software).

### Immunofluorescence staining on isolated myofibres and quantification

Skeletal myofibres from WT and *speg*-KO were isolated as previously described (Horstick et al., 2013). Briefly, embryos at each respective time point were digested with collagenase type II (LS004176, Worthington Biochemical Corporation) and plated on 12 mm circular coverslips. Samples were then fixed with 4% PFA for 20min at room temperature or 100% Methanol for 10 min at 4°C, permeabilized with 1X PBSTw (0.1% Tween 20 in PBS), incubated with blocking solution (0.2% TritonX-100, 0.2% BSA, 5% Goat Serum in PBS) for 30 min∼1 hr at room temperature, and then incubated with primary antibodies overnight at 4°C. Samples were stained in the dark with secondary antibodies for 1 hr at room temperature, and mounted with ProLong Gold Antifade Mountant (Invitrogen). Sample slides were dried at room temperature overnight in dark and stored in 4°C until imaging. Images were taken using a Leica SP8 Lightning Confocal microscope. The primary antibodies used were: rabbit polyclonal anti-SPEG (1:100; PA553875, Invitrogen), mouse monoclonal anti-RyR1 (1:100; 34C, DSHB), rabbit polyclonal anti-CACNA1S (1:100; ab203662, Abcam; which labels the DHPR protein), mouse monoclonal anti-SERCA1a (1:200; ab2819; Abcam), rabbit polyclonal anti-DES (1:100; D8281, Sigma), rabbit polyclonal anti-DNM2 (1:100; GTX127330; GeneTex), and mouse monoclonal anti-α-actinin (1:100; A7811; Sigma). The secondary antibodies used were anti-rabbit Alexa Fluor^®^ 488 (1:300, Invitrogen), and anti-mouse Alexa Flour^®^ 594 (1:300, Invitrogen).

To quantify immunofluorescence intensity, all Z-stack images were taken with the same microscope settings (i.e. objective, laser power, intensity, zoom, pixel size, and scanning speed). At least three fibres were imaged per group (WT or mutant) per myofibre prep immunostaining experiment, and three independent experiments were performed. For each image, average projection was generated using Fiji ImageJ, gray value was measured for signals (average of 3 measurements within the fibre) vs backgrounds (average of 3 measurements outside the fibre) using the square tool (area specified as 25 μm x 8 μm), and background value was subtracted from signal. The means of background-subtracted signals per group per experiment were plotted in GraphPad Prism version 8 (GraphPad Software), and compared using Student’s *t*-test.

### Transmission electron microscopy (TEM)

7 dpf larvae were anaesthetized using 0.1% tricaine and fixed in Karnovsky’s fixative [2.5% glutaraldehyde (GA) / 2% PFA in 0.1M cacodylate buffer, pH 7.5] at room temperature for 2 hrs, and re-fixed in fresh fixatives overnight at 4°C. The samples were then washed 3x 5 min in 0.1M cacodylate buffer (pH 7.5), post-fixed in 1% osmium (in 0.1M cacodylate buffer, pH 7.5) for 1.5 hr at room temperature, and washed 3x 5 min with 0.1M cacodylate buffer. Samples were then dehydrated with serial ethanol washes (70%, 90%, 95%, and 100%), infiltrated with Epon, and embedded in Epon to let polymerize in a 60°C oven for 24-48 hrs. Semi-thin (1 μm) and ultra-thin (90 nm) sections were cut using Leica Ultracut ultramicrotomes and transferred on 200 nm copper grids. Grids were post-stained with 2% uranyl acetate at room temperature for 20 min, washed 7x 1 min with MilliQ water, and stained with lead citrate for 5 min, followed by 7 x 1 min water wash. Samples were imaged using FEI Tecnai 20 transmission electron microscope.

### Measurement of electrically-evoked and caffeine-induced Ca^2+^ release in dissociated myofibres

For all experiments, individual myofibres were isolated by enzymatic digestions and then loaded with 5 µM fluo-4AM (Molecular Probes) for 45 min at room temperature in a normal rodent Ringer’s solution consisting of (in mM): 145 NaCl, 5 KCl, 2 CaCl_2_, 1 MgCl_2_, 10 HEPES, pH 7.4. Fibres were then transferred to dye-free rodent Ringer’s solution supplemented with 25 µm N-benzyl p-toluene sulfonamide (BTS) for 20 min at room temperature to block contractions. Fluo4-loaded fibres were excited at 480 ± 15 nm and fluorescence emission detected at 535 ± 20 nm was collected at 10 kHz using a photomultiplier system. Myoplasmic Ca**^2+^** transients in fluo4-loaded myofibres were stimulated by an electrical field stimulation protocol using a glass electrode filled with 200 mM NaCl placed adjacent to the cell of interest. The stimulation protocol consisted of five twitch stimuli delivered at 1 Hz followed by a single 5 sec, 10 Hz stimulation train, and then exposure to 10 mM caffeine for 30 sec in the absence of electrical stimulation. Peak change in fluo-4 fluorescence was measured and expressed as (F_max_-F_0_)/F_0_.

### Swimming assay and muscle performance quantification

To quantify muscle performance, 3 dpf and 5 dpf zebrafish were individually transferred to a 96-well plate and incubated in an optovin analog 6b8 (10 μM in 200 μL of embryo water, ChemBridge, Cat# 5707191) at 28.5°C for 5 min in the dark. Motor activity of the larvae was recorded and analyzed using ZebraBox (Viewpoint, France) as previously described (Zhao et al., 2019) with 30 sec light on, 1 min light off, 30 sec light on, 1 min light off, and 30 sec light on. Four independent experiments were conducted including WT vs *speg*-KO embryos, and n=24 larvae per group. Total distance travelled (mm) was plotted and analyzed using GraphPad Prism version 8 (GraphPad Software). Standard error of the mean (SEM) was calculated for each group, and one-way ANOVA was performed to test statistical significance.

### Protein extraction

Embryos were fin clipped at 3-4 dpf and genotyped using HRM analysis as described above. 5 dpf WT or *speg*-KO embryos were collected (∼20 embryos per group) and immediately stored at -80°C. Samples were homogenized using a Pellet Mixer (VWR, Cat# 47747-370) in 1X RIPA buffer (Cell Signaling, Cat# 9806) supplemented with Complete Mini EDTA-free Protease inhibitor tablets (46548400, Roche, ½ tablet per 5 mL lysis buffer) and phosphatase inhibitors (CA80501-130, VMR, 1:100). Lysates were chilled at 4°C for 10 min, sonicated and centrifuged at 12,000×*g* for 30 min at 4°C. Supernatants were collected, and protein concentration quantified using the Pierce™ BCA protein assay kit (Cat# 23225, Thermo zebrafisher Scientific).

### Western Blot

Protein lysates (40 µg/lane) were mixed with LDS sample buffer X4 (Invitrogen, B0007, X1) /DTT (100 mM) and boiled at 95°C for 5 min before loading. Samples were run at 100V, transferred using semi-dry transfer at 10V for 70min and resolved on PVDF membranes. Equal loading and transfer efficiency were assessed by total protein (REVERT^TM^ 700, Li-cor) staining prior to blocking. Membranes were blocked in 1X TBST containing 3% bovine serum albumin (BSA) for 1-2 hrs at room temperature, and then incubated overnight at 4°C with primary antibody in blocking solution. Membranes were washed and probed with secondary antibody (1:5000, anti-Rabbit-HRP, 1706515; anti-Mouse-HRP 1706516, BioRad) in blocking solution. Blots were imaged by chemiluminescence (Clarity Max™ ECL, BioRad) using the Gel Doc™ XR + Gel Documentation System (BioRad). Band signal intensities were determined using Fiji ImageJ software. All densitometry values are individually standardized to corresponding values of total protein stain and expressed as the fold difference from the average of the WT group of each blot. Primary antibodies used: anti-DES (1:1000; D8281, Sigma), anti-beta-actin (1:1000, 8226, Abcam), and Dnm2 (1:1000; GTX127330; GeneTex).

### Statistics

Post-capture analysis, including tests of statistical significance, was performed using Microsoft Excel 2016 (Microsoft) and GraphPad Prism version 8 (GraphPad Software). The difference between three or more groups was assessed by ordinary one-way ANOVA followed by Tukey’s multiple comparisons test. The difference between two groups was assessed by 2-tailed Student’s *t*-test or by 2-tailed Mann-Whitney test. All survival curves were assessed by Mantel-Cox test. Differences were considered to be statistically significant if \**P* < 0.05, \*\**P* < 0.01, \*\*\**P* < 0.001, or \*\*\*\**P* < 0.0001. All data unless otherwise specified are presented as mean ± SEM.

---

**Supplementary Figure 1.**
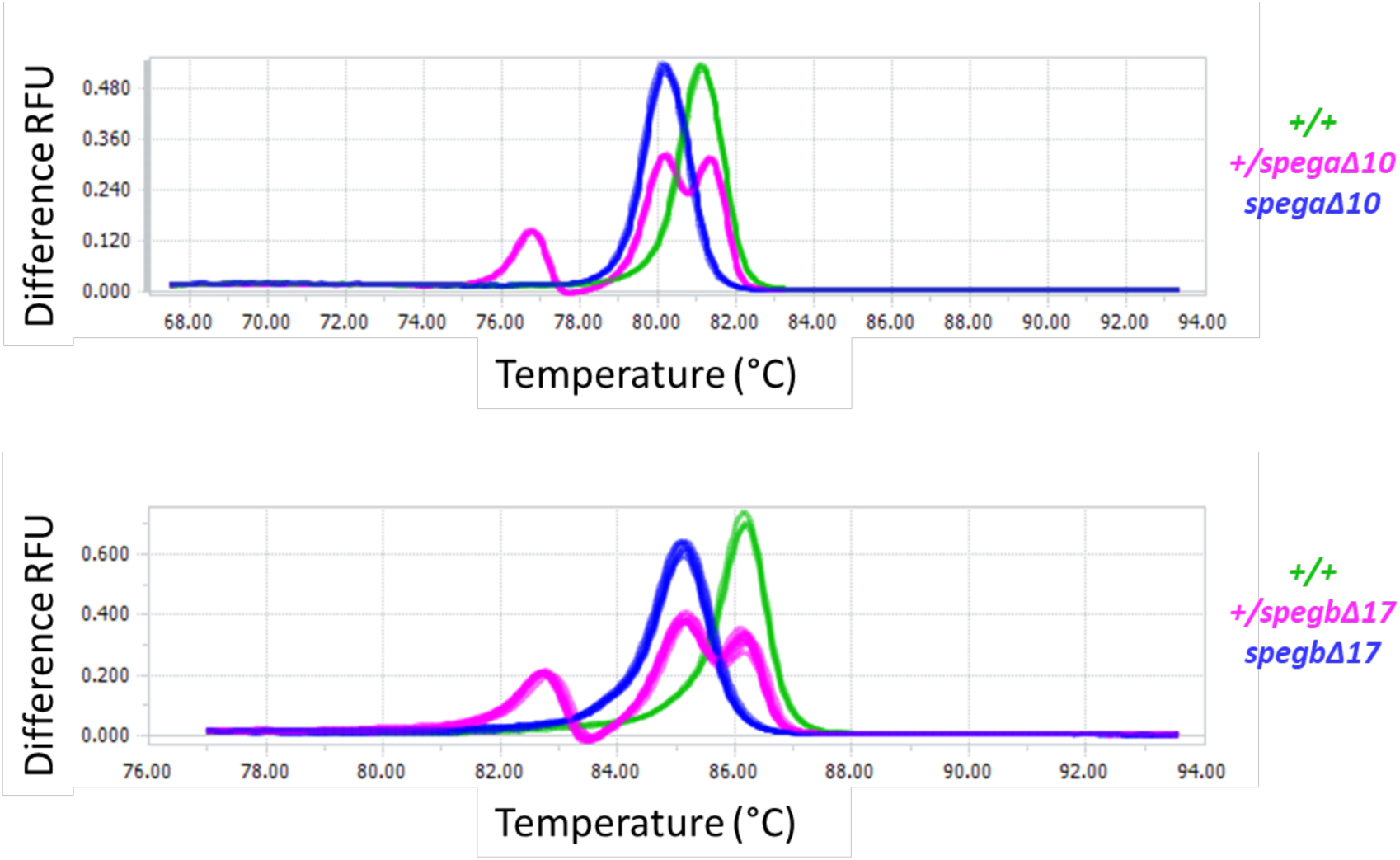
*spegaΔ10;spegbΔ17* mutant zebrafish genotyping. High Resolution Melting (HRM) analysis was used to genotype *spega* (top) and/or *spegb* (bottom) CRISPR lines. Wildtype (*green*), Heterozygous (*magenta*), and Homozygous mutants (*blue*). RFU: relative fluorescence units.

**Supplementary Figure 2.**
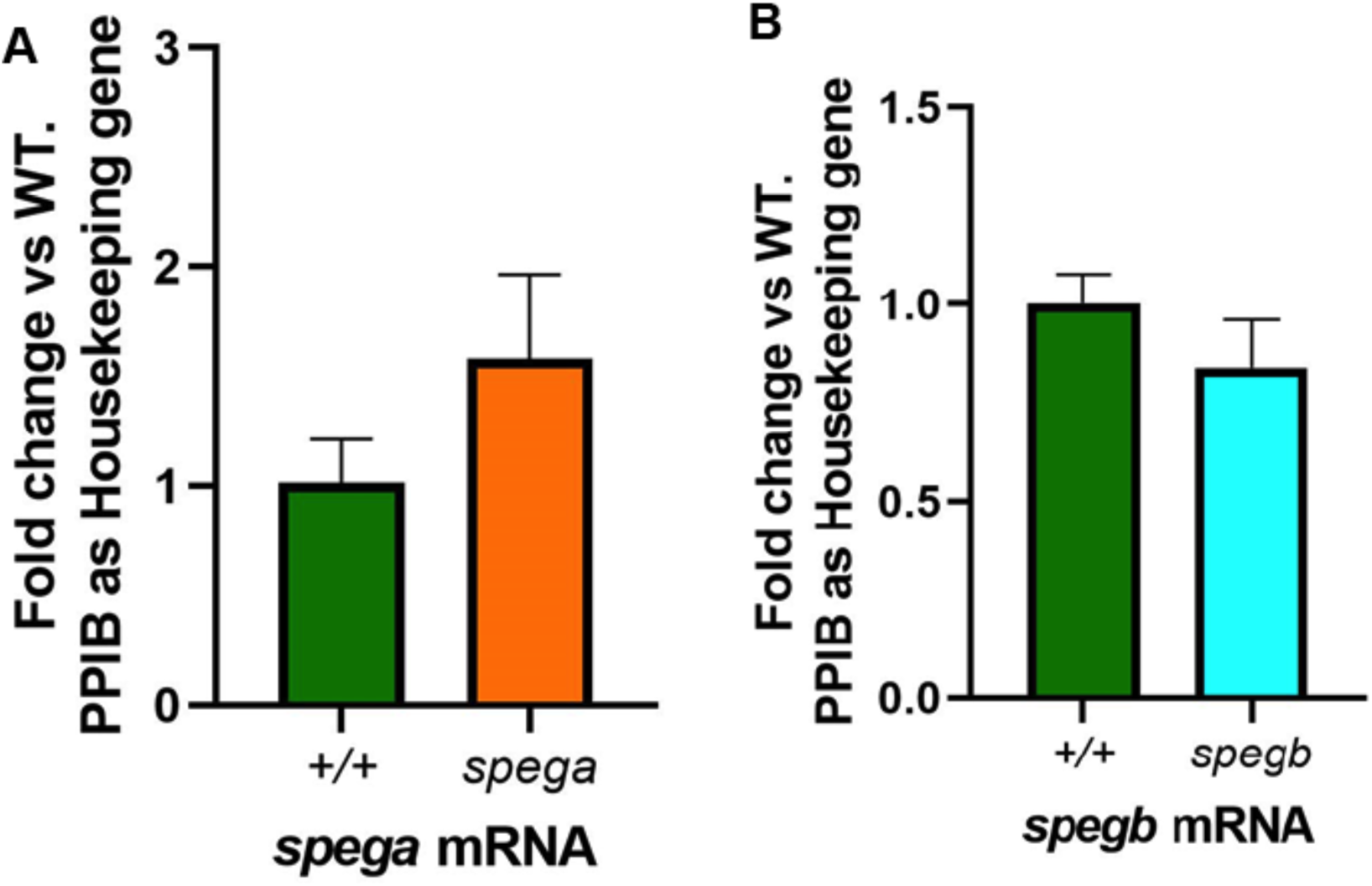
RT-qPCR analysis on single mutants. No significant changes were observed at 7 dpf for the levels of *spega* mRNA in *spegaΔ10*-KO (**A**), or *spegb* mRNA in *spegbΔ17*-KO (**B**). Levels of expression were first normalized to the housekeeping gene *ppib*, and then to WT. Results represent three independent experiments (technical triplicates per experiment). N=10-15 embryos per genotype per independent experiment. Columns and error bars represent Mean ± SEM. Two-tailed Student’s *t*-test was performed: ns, not significant.

**Supplementary Figure 3.**
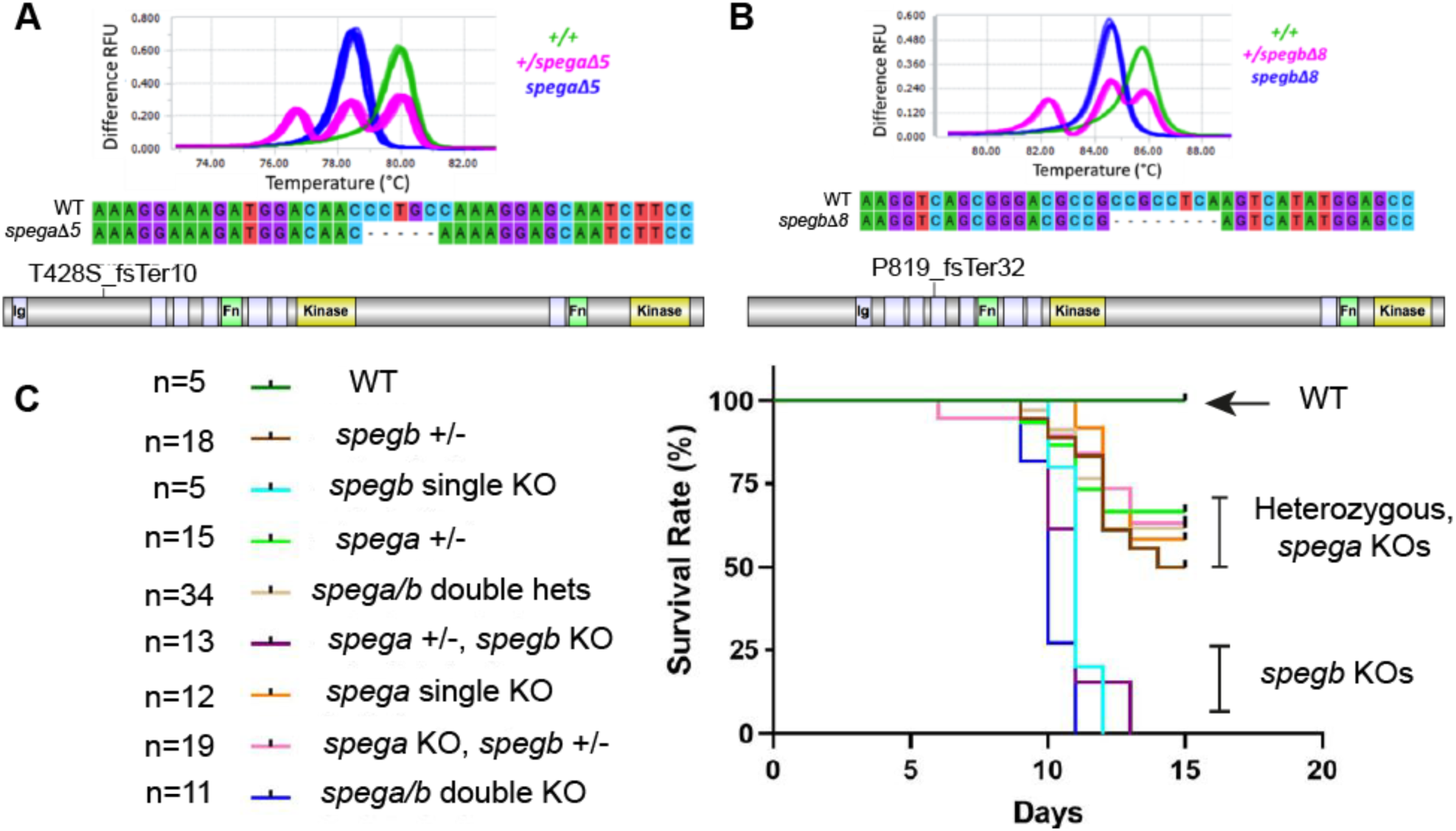
Reduced survival in *spegaΔ5;spegbΔ8*. Additional mutant lines were generated to validate the *speg*-DKO (*spegaΔ10;spegbΔ17)* survival phenotype. **(A** and **B)** HRM and Sanger sequencing confirmed the genotypes of (**A**) *spegaΔ*5, which carries a 5-bp deletion in exon 5 of *spega*, and (**B**) *spegbΔ8,* which carries an 8-bp deletion in exon 8 of *spegb.* Both are predicted to cause frame shift (fs) and premature stop codons. (**C**) Survival analysis across all *spegaΔ5* and/or *spegbΔ8* lines: survival was reduced in *spegb* KOs, *spegb* KO/*spega* +/-, and *spega/b* double knockouts, with a median survival of 11, 11 and 10 dpf, respectively, and a maximum survival of 12, 13 and 11 dpf, respectively. This represents a significant decrease in survival as zebrafish can live up to 1.5-2 years (****P<0.0001), Mantel-Cox test. In contrast, WT and *spega* KOs have similar lifespans.

**Supplementary Figure 4.**
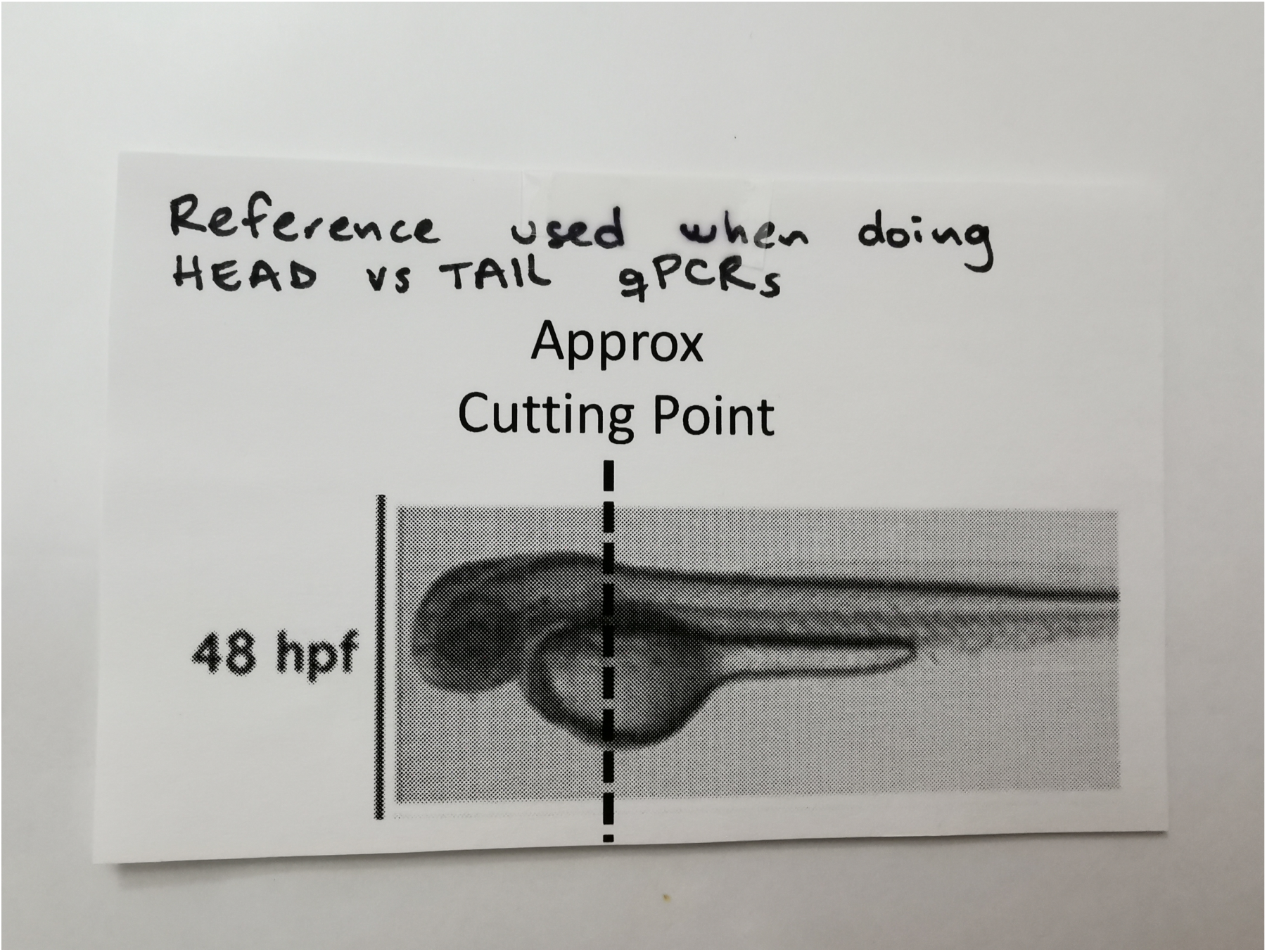
An illustration showing the approximate cutting point (dotted line) to separate head vs tail for RT-qPCR analyses.

